# MicroRNA-423-5p Mediates Cocaine-Induced Smooth Muscle Cell Contraction by Targeting Cacna2d2

**DOI:** 10.1101/2023.02.08.527687

**Authors:** Derek M Dykxhoorn, Huilan Wang, Andrea Da Fonseca Ferreira, Jianqin Wei, Chunming Dong

## Abstract

**Background:** Cocaine abuse increases the risk of atherosclerotic cardiovascular disease (CVD) and causes acute coronary syndromes (ACS) and hypertension (HTN). Significant research has explored the role of the sympathetic nervous system mediating the cocaine effects on the cardiovascular (CV) system. However, the response of the sympathetic nervous system alone is insufficient to completely account for the CV consequences seen in cocaine users. Here, we examined the role of microRNAs (miRNAs) in mediating the effect of cocaine on the CV system. MiRNAs regulate many important biological processes and have been associated with both response to cocaine and CV disease development. Multiple miRNAs have altered expression in the CV system (CVS) upon cocaine exposure. Herein, we examined the role of microRNA-423-5p and the downstream signaling events in regulating cocaine-induced mouse aortic smooth muscle cell (SMC) contraction.

**Methods:** To understand the molecular mechanisms underlying the cocaine response in the CV system, we studied the role of miRNA-423-5p and its target Cacna2d2 in the regulation of intracellular calcium concentration and SMC contractility, a critical factor in the modulation of blood pressure (BP). We used in vivo models to evaluate BP and aortic stiffness.

**Results:** *In vitro*, Cocaine treatment decreased miR-423-5p expression and increased Cacna2d2 expression, which led to elevated intracellular calcium concentrations and increased SMC contractility. Overexpression of miR-423-5p, silencing of its target Cacna2d2, and treatment with a calcium channel blocker reversed the elevated SMC contractility caused by cocaine. In contrast, suppression of miR-423-5p increased the intracellular calcium concentration and SMC contractibility. *In vivo*, overexpression of miR-423-5p ameliorated the increase in BP and aortic stiffness associated with cocaine use.

**Conclusions:** MiR-423-5p regulates SMC contraction by modulating Cacna2d2 expression increasing intracellular calcium concentrations. Modulation of miR-423-5p—Cacna2d2—Calcium transport pathway may represent a novel therapeutic strategy to improve cocaine-induced hypertension and aortic stiffness.

## INTRODUCTION

Cocaine is a powerful sympathomimetic agent derived from the leaves of the *Erythroxylum coca* plant. With the exception of marijuana, cocaine remains the most commonly used drug of abuse and is the leading cause of drug abuse-related emergency room visits^1^. Cocaine is associated with a wide range of cardiovascular (CV) complications, including acute coronary syndromes, heart failure, cardiomyopathies, arrhythmias, stroke, hypertension (HTN), and aortic dissection^2-8^. Cocaine has multiple CV and prothrombotic effects that may contribute to the development of CV disease. Cocaine consumption causes a dose-dependent increase in blood pressure (BP)^9^ and heart rate^10,11^. In addition, cocaine induces coronary and peripheral arterial vasoconstriction ^12-15^. Cocaine stimulates the sympathetic nervous system by blocking the reuptake of catecholamines at the presynaptic adrenergic terminals leading to the accumulation of catecholamines at the post-synaptic receptors and increasing the sensitivity of adrenergic nerve endings to norepinephrine (NE)^12,16^. This was thought to be the mechanism underlying the cocaine effects on the CV system. However, when cocaine or cocaine methiodide (CM, which does not enter the CNS) was administered 5 minutes before giving a calcium channel blocker Nifedipine, neither cocaine nor CM changed the effect of NE on heart rate and blood pressure (BP)^17^. This observation suggests that the NE potentiation mechanism cannot fully explain cocaine’s effects on the CV system. Despite the frequency of CV complications in individuals using cocaine, few, if any, studies have explored the molecular mechanisms underlying the effects of cocaine on the CV system beyond its sympathomimetic effect. Although some studies implicated intracellular calcium overload^18^, oxidative stress^19,20^, and mitochondrial dysfunction^21,22^ in cocaine-induced cardiotoxicity, these mechanisms were all studied in the context of the effects of catecholamines.

Micro (mi) RNAs are small non-coding RNAs that post-transcriptionally regulate gene expression by binding to the 3’ UTR of target transcripts leading to translational repression and/or mRNA degradation^23-25^. MiRNAs have been shown to play important roles in CV development and disease, vascular aging, and response to drug exposure^26-30^. The role of miRNAs in mediating the effects of cocaine exposure on the central nervous system has been examined. For example, miR-212 has been shown to play a prominent role in the vulnerability to cocaine addiction by controlling two complementary mechanisms – amplification of CREB signaling^31^ and reduction of MeCP2/BDNF transmission in the striatum^32^. Additional miRNAs have been found to be dysregulated in various regions of the brain following cocaine exposure (reviewed in^33^). However, little is known about the role of miRNA pathways in mediating the cocaine effects in the CV system. Recently, we reported that miR-30c-5p was upregulated in the aortas of mice treated with cocaine. MiR-30c-5p directly targeted the redox molecule malic enzyme 1 (Me1) leading to increased reactive oxygen species (ROS) levels. Interestingly, the silencing of miR-30c-5p in mouse aortas abrogated the cocaine-induced increase in ROS, resulting in partial normalization of BP and aortic stiffness^30^.

To investigate the potential involvement of additional miRNA—mRNA pathways in mediating the cocaine effects in the CV system, we examined the expression of miR-423-5p in aortas from a mouse model of cocaine use/abuse. MiR-423-5p has previously been linked to CV disease and shown to induce apoptosis in cardiomyocytes by targeting O-GlcNAc transferase^34^, as well as, being proposed as a potential biomarker for CV disease, in particular heart failure ^35-37^. In this study, miR-423-5p was shown to be downregulated in cocaine exposed compared to the control treated mouse aortas. *In silico* analysis identified *Cacna2d2* –- the gene encoding the α2δ-2 subunit of voltage-dependent calcium channels - as a putative target of miR-423-5p. Luciferase reporter assays confirmed that *Cacna2d2* was a direct target of miR-423-5p. Silencing of Cacna2d2 – either through the overexpression of miR-423-5p or a Cacna2d2 siRNA – led to decreased contractility of smooth muscle cells (SMCs) *in vitro*. Importantly, *in vivo* overexpression of miR-423-5p in mouse SMCs reduced cocaine-induced elevation of BP and aortic stiffness. These results support an important role for the miR-423-5p – Cacna2d2 axis, in addition to our previously characterized miR-30c-5p – ME1 – ROS pathway, in the regulation of vasoconstriction resulting from cocaine exposure.

## METHODS

### Animals

Male C57BL/6 mice aged 8-10 weeks were purchased from Charles River Laboratory (Hollister, CA). Animals were treated according to National Institute of Health guidelines. Mouse protocols were approved by the Animal Care and Use Committee (IACUC) of the University of Miami Miller School of Medicine. Mice received intraperitoneal (I.P.) injections of cocaine, CM (20mg/kg, NIDA Drug Supplied) or Saline for 10 consecutive days. After the last injection, mice were measured for PWV and euthanized. Aortas were collected. In parallel animal experiments, mice received tail vein injection of lentiviruses encoding miR-423-5p or empty vector control (both from Biosettia, CA) driven by SMC-specific promotor *SM22*α at the dose of 3×10^5^ IU per mouse daily, every other day injection for a total of 3 injections over the period of 5 days. Then mice received daily injection of cocaine or saline for 10 consecutive days as described above.

### Blood Pressure and Pulse Wave Velocity Measurement

BP and PWV were measured as previously described^30^. Briefly, CODA BP monitor (Kent Scientific, Connecticut) was used to measure mouse systolic and diastolic BP followed the manufacturer’s instructions. BP was measured before the first injection of cocaine (baseline BP), then at days 1, 3, 5, 7 and 10 one hour after the cocaine injection. PWV was measured before the first injection and at the end of the last injection. Mice were anesthetized with 2% isoflurane and laid on a platform. Blood flow velocity was measured at the middle level of the ascending aorta. By placing a 420-40 MHZ Doppler probe to the right of the upper sternum, the aortic arch velocity signal was obtained. Three measurements were recorded for each mouse.

### Lentivirus Production and SMC Cell Lines

C57BL/6 mouse primary aortic SMCs were purchased from Cell Biologics (Cell Biologics, Chicago, IL). Cells were cultured to passage 4 in the Complete Smooth Muscle Cell Medium (Cell Biologics), in the culture plates coated with Gelatin (Cell biologics). The lentiviral vectors for the scrambled miR-Ctrl, miR-423-5p, miRZip-Ctrl, and miRZip423-5p (anti-miR-423-5p) were purchased from System Biosciences Inc (Palo Alto, CA). Lentiviral vectors encoding miR-Ctrl, miR-423-5p, miRZip-Ctrl, miRZip-423-5p were generated by co-transfection of the specific lentiviral vector with the lentiviral packaging plasmid, pCMV-D8.2, and the Vesicular stomatitis virus envelope glycoprotein (VSVg) expression construct, pCMV-VSV-G, at a ratio of 3:2:1 in HEK 293T cells (Sigma Aldrich) using Lipofectamine 2000 (Thermo Fisher Scientific). 48 hours after transfection, culture medium was collected filtered and concentrated using the Lenti-X™ Concentrator (Takara). Viral particles were suspended in fresh DMEM medium and stored at - 80°C. ShRNA for Cacna2d2 and the control shRNA lentiviral particles were purchased from Santa Cruz Biotechnology. To establish stable SMC cell lines, SMCs were transduced with specific lentiviral vectors. Briefly, SMCs were seeded at the density of 2×10^5^ per well in 6-well plates. After 2 hours, viral vectors were added to the cell culture medium together with 5μg/ml polybrene (Millipore Sigma). The stably transduced cells were selected by treating the cells with 10 μg/ml puromycin.

### Site-Directed Mutagenesis in Cacna2d2 and Luciferase reporter assays

Full length wild Cacna2d2-3’UTR was cloned downstream of a Gaussia luciferase reporter gene in the pEZX-MT05 vector (GeneCopoeia). Site-directed mutagenesis was used to introduce mutations into the putative miR-423-5p binding sites on the Cacna2d2 3’UTR using the QuickChange II XL site-direct mutagenesis kit (Agilent) according to the manufacturer’s protocol. The pEZX-MT05 vector contains a constitutively expressed secreted alkaline phosphatase (SeAP) reporter gene, which served as an internal control for transfection normalization. Luciferase reporter assays were performed by co-transfection of miR-423-5p or miR-Ctrl construct (System Biosciences) with either the wild type Cacna2d2-3’UTR or the Cacna2d2 3’UTR bearing mutations in the miR-423-5p binding sites (Cacna2d2 mut 3’UTR) in HEK 293 cells (Millipore Sigma). Briefly, HEK 293 cells were seeded at 32,000 cells/well in 96-well plates and cultured overnight. The following day the miR-423-5p or miR-Ctrl vector was cotransfected with either the Cacna2d2-3’UTR or the Cacna2d2 mut 3’UTR reporter vectors using Lipofectamine 2000 (Thermo Fisher Scientific) according to the manufacturer’s protocol. Luciferase and alkaline phosphatase activities were assayed using the Secrete-Pair™ Dual Luminescence assay kit (Genecopoeia) and read on the Centro XS3 LB960 Microplate Luminometer (Berthold Technologies, Germany).

### Real–Time quantitative PCR (RT-PCR) for mRNAs and miRNAs

Total RNA containing miRNAs was extracted from untransfected SMCs and cells transduced with miR-423-5p, miRZip-423-5p, scramble miR-Ctrl, miRZip-Ctrl, shRNA-Ctrl, shRNA-Cacna2d2 using the miRNeasy Mini Kit (Qiagen, Germantown, MD) according to the manufacturer’s instructions. RNA concentrations were assessed using the NanoDrop™ 2000 Spectrophotometer (Thermo Fisher Scientific). Complementary DNA was produced using the High-Capacity cDNA Reverse Transcription kit (ABI) according to the manufacturer’s protocol. Quantitative real time PCR (qRT-PCR) analysis was performed using the IQ SYBR Green Supermix (Bio-Rad) according to the manufacturer’s protocol and read on the IQ5 multicolor Real-Time PCR Detection system (Bio-Rad). The Primers for Cacna2d2 were: Cacna2d2 forward primer: 5’-ccgctcttgctcttgctg-3’; Cacna2d2 reverse primer: 5’-ccagtgctgcatcgtgtg-3’. GAPDH was used as an internal control. Quantitation of miR-423-5p expression levels was assessed using the Taqman™ MicroRNA Reverse Transcription kit (Thermo Fisher Scientific) and the specific Taqman™ MicroRNA Assay kit and the Universal PCR Master Mix No AmpErase UNG (both Thermo Fisher Scientific). The RT-PCR reaction were performed on ABI 7900HT Fast Real-Time PCR system. The U6 small nucleolar RNA was used as the housekeeping small RNA reference gene for normalization of sample input.

### Cytosolic free calcium measurement

The intracellular free Ca^2+^ fluorescence images were obtained by free Ca^2+^ sensitive Fluo3-AM green fluorescence staining and the intracellular free Ca^2+^ concentrations were measured by FACS. The Fluo-3AM (Thermo Fisher Scientific) stock solution was prepared by first dissolving 50μg of Fluo-3-AM dye in 35μl DMSO. 14 μl of 10% Pluronic F-127 (Thermo Fisher Scientific) was added to the dissolved Fluo-3-AM solution and then mixed with 117μl of PBS. Cells were cultured and treated with Cocaine (150μm) or CM (150μm) for 48 hours, then washed and stained with Fluo3-AM solution for 40 min at 37°C. Fluorescence images were obtained using the EVOS FL Auto Cell Imaging System (Thermo Fisher Scientific). For FACS analysis of intracellular free Ca^2+^, cells were cultured and treated in the same way as Fluo3-AM staining. Briefly, cells were collected and washed with 1 x HBSS containing 1% fetal bovine serum then suspended in 1.0 ml of 1 x HBSS containing 2.5μg/ml of Fluo-3-AM dye and 2.5μm of the anion carrier inhibitor probenecid (Thermo Fisher Scientific) and incubated for 45 min at room temperature on an orbital shaker in the dark, washed and resuspended in 1 x HBSS containing probenecid, and incubated for 20 min in the dark. Fluorescence intensity in the stained cells was measured by FACS analysis. Ionomycin (Millipore Sigma) was used as a positive control and was added to the suspended cells 10 min before imaging and FACS analysis.

### Collagen gel contraction assay

The ability of SMCs to contract in response to CM, Cocaine, or the Calcium channel blocker Nimodipine (NM) was evaluated by using CytoSelect™ 24-Well Cell Contraction Assay Kit (Cell Biolabs) according to the manufacturer’s protocol. Briefly, SMCs were harvested and suspended in cell culture medium at 1 × 10^6^ cells/ml. Collagen gel working solution was made according to the manufacturer’s instructions. Two parts of suspension cells were mixed with eight parts of collagen gel working solution to make cell contraction matrix and 0.5 ml of the cell contraction matrix was added to each well of the 24-well cell contraction plate. Collagen gels were allowed to solidify at 37°C in 5% CO_2_ for 1 hour. After the collagen gel solidified, 1.0 ml of culture medium containing 150 μM of Cocaine, 150 μM of CM, or ET-1 was added atop the collagen lattice. Following the 48h treatment, the gel is gently released from the sides of the culture plate and the contractility of the matrix is measured by digital imaging. The contractibility of SMCs infected with lentiviruses containing miR-Ctrl, miR-423-5p, miRZip-423-5p, shCtrl, shCacna2d2 was evaluated and compared. All experiments were performed in triplicates.

### Immunofluorescence staining

Immunofluorescence staining was performed on the frozen aortic sections. Briefly, mice were sacrificed and the aortas were flash frozen. Slides were obtained by cutting blocks in CM1850 microtome (Leica Biosystems). Frozen sections were fixed in ice cold 4% paraformaldehyde for 10 min at 4°C, permeabilized in 0.3% Triton X-100/PBS for 5 min and incubated in 5% BSA for 30 min at room temperature to facilitate the blocking of nonspecific binding. Subsequently slides were incubated with primary Cacna2d2 antibody at 1:50 dilation (Biorbyt) or isotype control at 4°C overnight. Slides were washed with PBS and incubated with goat anti-rabbit AlexaFluor 488 diluted 1:300 (Molecular Probes, Grand Island, NY) and Hoechst 33342 dilution of 1:300 (Molecular Probes, Grand Island, NY) for 60 min, washed, and mounted with Permount™ mounting media (Fisher Scientific). Images were acquired using the EVOS FL Auto Cell Imaging System.

### Statistical analysis

All results are expressed as mean ± standard deviation (SD). Student’s two tailed *t*-test was used for all statistical analysis. Statistical analysis was performed by using SPSS16.0 computer software. A *p*-value <0.05 was considered as statistically significant.

## RESULTS

### 1. Cocaine and cocaine methiodide (CM) exposure increases BP, aortic stiffness, and alters miR-423-5p and Cacna2d2 expression in the mouse aort

Cocaine has been linked to increased risk for CV disease ^9, 38-40^. To assess the impact that cocaine exposure has on miRNA expression in the aorta, we developed a model of cocaine use/abuse in C57BL/6 mice ^30^. In this model, 8-10 week old mice were given daily intraperitoneal injections of cocaine, cocaine methiodide (CM), or saline. Compared to the saline treated mice, both systolic and diastolic BP was significantly increased in the cocaine and CM treated animals throughout the treatment course as previously described^30^. In addition, the cocaine and CM treated mice showed increased aortic stiffness, as measured by pulse wave velocity (PWV) on day 0 and then two days following the final injection of cocaine^30^. We had previously shown that cocaine exposure results in an increase in miR-30c-5p levels leading to decreased expression of Malic enzyme-1 (Me-1) and increased reactive oxygen species. Reducing miR-30c-5p expression or treating mice with the free oxygen radical scavenger with N-acetyl cysteine (NAC) led to reduced pulse wave velocity – a measure of aortic stiffness^30^. Quantitative real time PCR (qRT-PCR) analysis also showed that miR-423-5p expression was decreased in the aortas of cocaine- and CM-treated mice compared to the control (saline) treatment (**Fig 1A**). *In silico* analysis of miRNA targets using prediction programs – TargetScan and miRdb – identified Cacna2d2 as a putative miR-423-5p target^41,42^. Furthermore, the expression of Cacna2d2 was elevated in cocaine- and CM-treated compared to the control (saline) treated mouse aortas (**Fig. 1A**). Immunohistochemical staining of the mouse aortas confirmed the increase in Cacna2d2 expression in response to CM and cocaine treatment, relative to saline control (**Fig 1B & C**). To determine if the anticorrelation of miR-423-5p with Cacna2d2 expression was due to direct inhibition of Cacna2d2 by miR-423-5p, luciferase reporter assays were performed. The 3’UTR of Cacna2d2 was cloned into a luciferase reporter construct which was transfected into HEK293T cells along with either miR-423-5p or a control miRNA (miR-Ctrl). The cotransfection of miR-423-5p with the Cacna2d2 3’UTR construct showed a significant decrease in luciferase activity compared to miR-Ctrl treatment (**Fig 1 D and E**). Three potential miR-423-5p binding sites – one highly conserved site and two with lower binding properties – have been identified *in silico* in the 3’ UTR of Cacna2d2 (**Fig 1D**). Site-directed mutagenesis was used to introduce mutations that should disrupt the interaction between miR-423-5p and the three potential miR-423-5p binding sites Cacna2d2 3’ UTR. Similar to the miR-Ctrl treatment, the co-transfection of miR-423-5p with the mutated version of Cacna2d2 3’ UTR (mtCacna2d2 3’ UTR) was unable to silence luciferase expression (**Fig 1E**). These results show that Cacna2d2 is a direct target of miR-423-5p.

**Figure 1.**
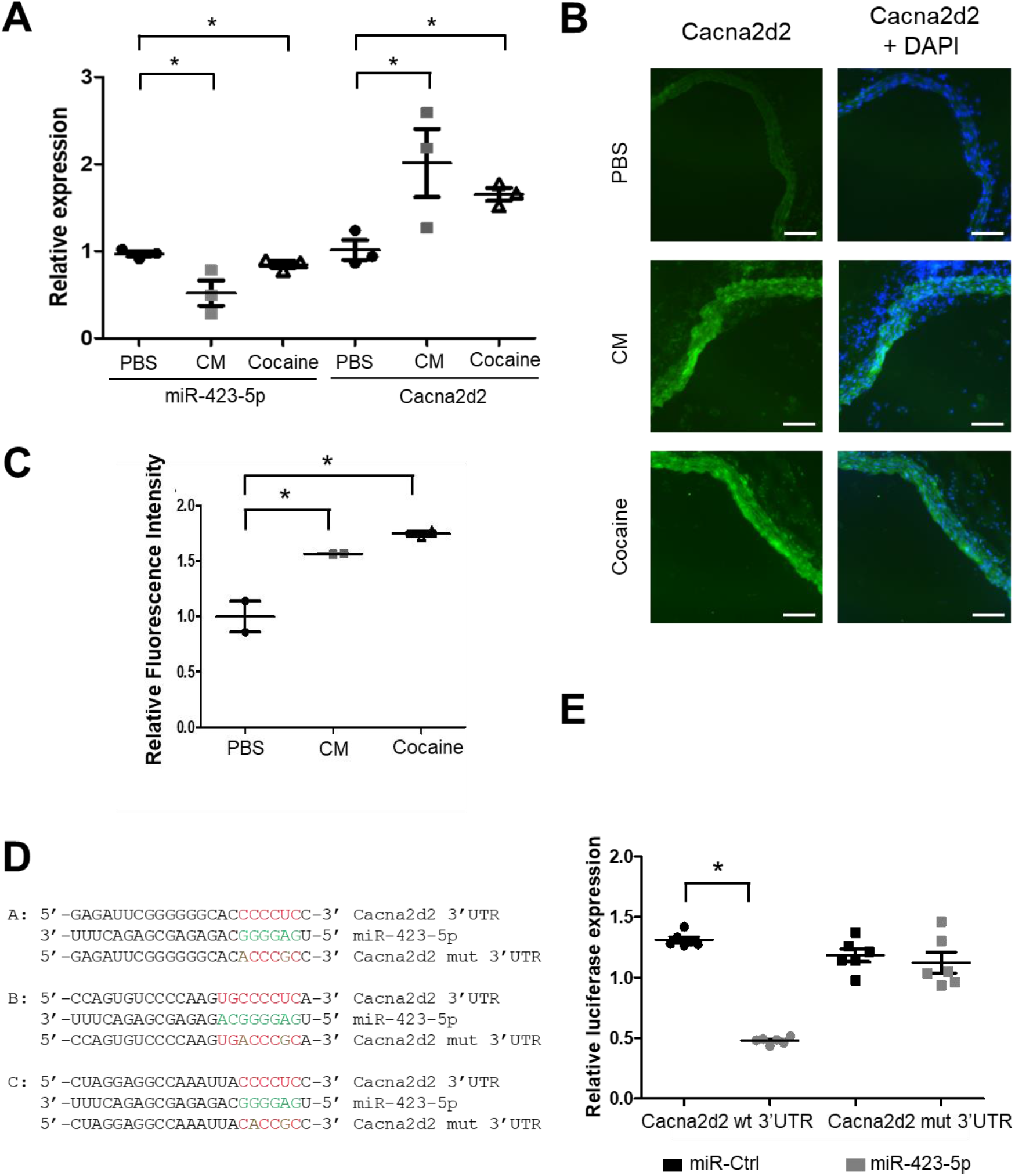
Cocaine and CM suppress miR-423-5P expression and increase Cacna2d2 expression in the mouse aortas. (**A**) QRT-PCR analysis was performed using RNA extracted from aortas isolated from mice treated with saline, cocaine, or CM for 10 consecutive days. Cocaine and CM resulted in decreased miR-423-5p expression and increased Cacna2d2 expression compared to saline treatment (\**p*<0.05 vs. saline). (**B**) Immunohistochemical (IHC) analysis of Cacna2d2 levels in aortic sections showed cocaine and CM treatment induced increased levels of Cacna2d2 protein expression compared to saline. (**C**) The intensity of Cacna2d2 positive staining was quantified using Image J software (* p<0.05 vs. saline). (**D**) *In silico* analysis identified 3 putative miR-423-5p binding sites in the Cacna2d2 3’ UTR. Point mutations were introduced in these three sites to determine the potential binding of miR-423-5p to the Cacna2d2 3’UTR. (**E**) Luciferase reporter assays were performed by co-transfection of HEK-293 cells with luciferase reporter constructs containing the WT or mutant Cacna2d2 3’ UTR and either the miR-423-5p or a non-specific control miRNA (miR-Ctr) expression vector. Gaussia luciferase activities (GLuc), measured and normalized to secreted alkaline phosphatase (SeAP) activities, show that Cacna2d2 is a direct target of miR-423-5p. (*p<0.05 vs. miR-Ctrl).

### 2. Cocaine and CM treatment increase intracellular free calcium [Ca^2+^] and induce primary mouse aortic SMCs contraction

The mice in the cocaine abuse/use model showed elevated BP and aortic stiffness ^30^. Cocaine abuse has been shown to induce vasoconstriction ^43-47^. Ca^2+^ plays a central role in excitation-contraction coupling in vascular SMCs among other functions (reviewed in^48^). Therefore, the regulation of intracellular Ca^2+^ concentration ([Ca^2+^]_i_) is crucial for the proper functioning of SMCs. Cacna2d2 encodes a subunit of the L-type Ca_v_1.2 channels that are key regulators of Ca^2+^ influx and myogenic tone^49-51^. To examine whether cocaine treatment caused the [Ca^2+^]_i_ changes in primary mouse aortic SMCs, intracellular Ca^2+^ levels were measured in mouse SMCs in the presence or absence of cocaine or CM. SMCs were cultured and treated with saline, cocaine, CM, or the calcium ionophore ionomycin, and the cells were stained with the Ca^2+^-specific fluorescent dye, Fluo-3 AM (**Fig 2A and B**). Fluorescence imaging of cocaine or CM treated cells showed an increase in intracellular Ca^2+^ compared to the saline control treated cells (**Fig 2A**). Flow cytometric analysis of Fluo-3 AM stained cells confirmed the increase in intracellular Ca^2+^ following treatment with cocaine or CM (**Fig 2B**). The level of fluorescence seen in response to cocaine or CM treatment was similar to the level observed when the cells were treated with the ionophore ionomycin (positive control). These results show that treatment with cocaine and CM led to an increase in [Ca^2+^]_i_ levels in SMCs.

**Figure 2.**
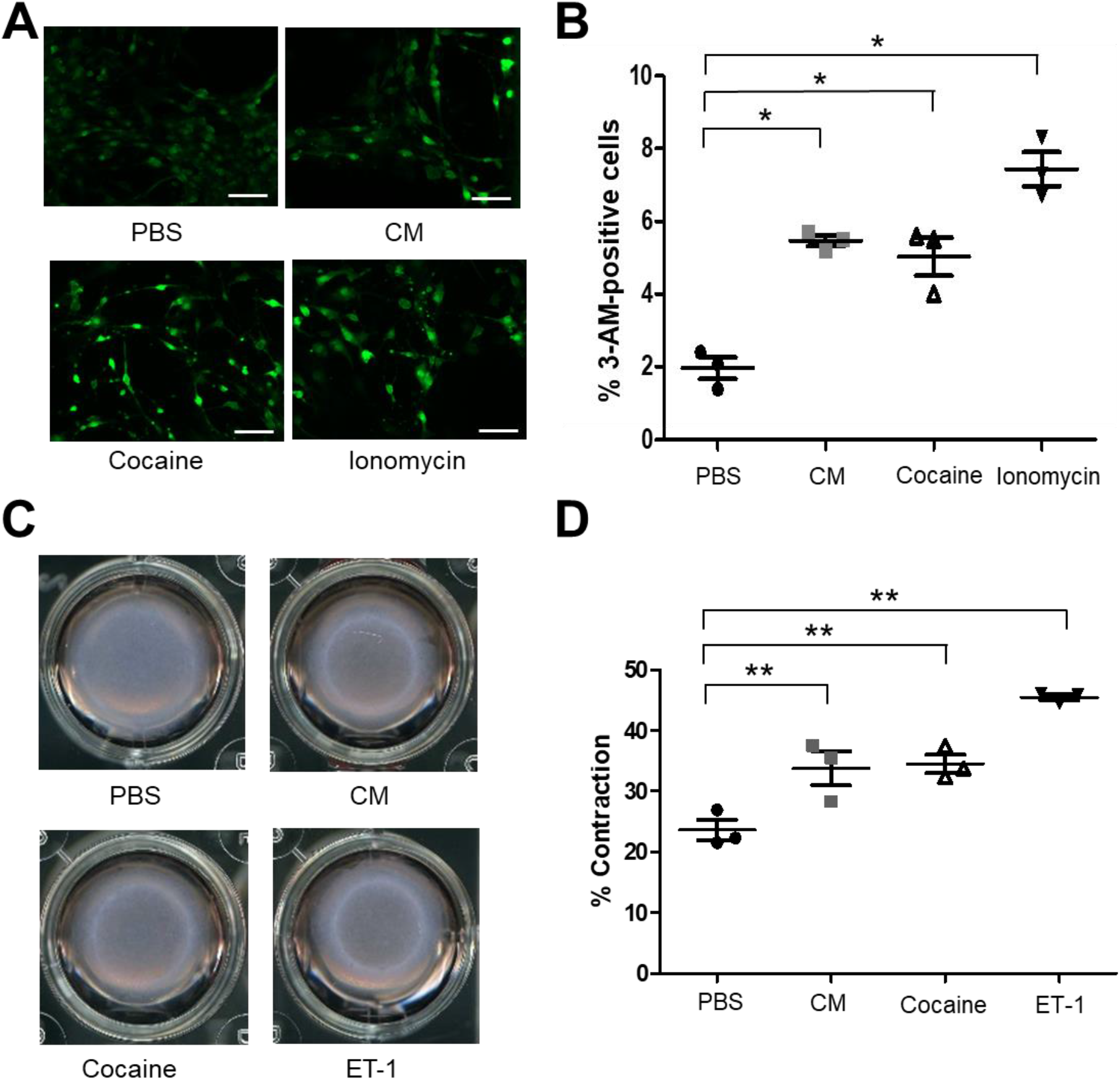
Cocaine and CM increase intracellular free calcium [Ca^2+^] and induce primary mouse aortic SMC contraction. (**A**) Fluorescence image analysis of free cytosolic Ca^2+^ in mouse aortic SMCs treated with PBS, CM, cocaine or the calcium ionophore ionomycin (positive control) and stained with the calcium specific dye Fluo-3 AM show that cocaine and CM increase the level of intracellular free calcium in SMCs. (**B**) Quantification by Image J software analysis fluorescence positive cells (*p<0.05 vs. PBS). (**C**) Mouse Aortic SMCs were exposed to saline, cocaine, CM or the vasoactive peptide ET-1 (positive control), and their contractility was measured using the collagen gel contraction assay. Representative images of SMC contractility in the collagen contraction assay show increased SMC contraction in response to cocaine and CM. (**D**) Quantification of collagen gel contraction show both cocaine and CM substantially increase SMC contraction relative PBS (*p<0.05 vs PBS).

To test the effect of cocaine and CM on the contractility of SMCs, a cell contraction assay was performed by embedding the SMCs in a collagen matrix that was attached to the well of the tissue culture plate. The initial period of attachment to the tissue culture plate allows for mechanical loading and stress fiber formation. 48 hours following plating, the collagen matrices are gently released from the tissue culture plate resulting in mechanical unloading and contraction of the cell-embedded collagen matrix and the contraction index measured. Treatment with cocaine or CM resulted in increased contraction of the collagen matrix compared to the PBS treated cells (**Fig 2C and D**). **Figure 2C** shows bright field images of the matrix embedded with SMCs and treated with saline, cocaine, CM, and endothelin-1 (potent endogenous vasoconstrictor molecule). The contractile index was quantified for each treatment (**Fig 2 D**). Treatment with cocaine or CM increased SMC contraction compared to the saline treatment. This increase in contractility was similar to that observed with endothelin-1 treatment (positive control for SMC contraction). These results show that cocaine and CM increase the [Ca^2+^]_i_ and contractility of SMCs compared to PBS treated cells.

### 3. Modulation of the miR-423— Cacna2d2 axis alters intracellular free calcium [Ca2+]_i_ and contractility of mouse aortic SMCs

Treatment with cocaine or CM increased [Ca^2+^]_i_ levels and the contractility of SMCs. We sought to determine if cocaine/CM were exerting their effect on SMC contractility through the miR-423-5p – Cacna2d2 pathway. To that end, the levels of miR-423-5p and Cacna2d2 were modulated in SMCs, and these cells were assayed for the impact of these changes on SMC contractility. As expected, overexpression of miR-423-5p in SMCs led to a significant decrease in Cacna2d2 expression (**Fig 3 A-B**). This decrease in Cacna2d2 expression was seen even in cells that were exposed to cocaine **(Fig 3B)**. Conversely, the silencing of miR-423-5p using the miR-423-5p antagomir, miR-Zip-423-5p, led to an increase in Cacna2d2 levels both in the absence and in the presence of cocaine **(Fig 3B)**.

**Figure 3.**
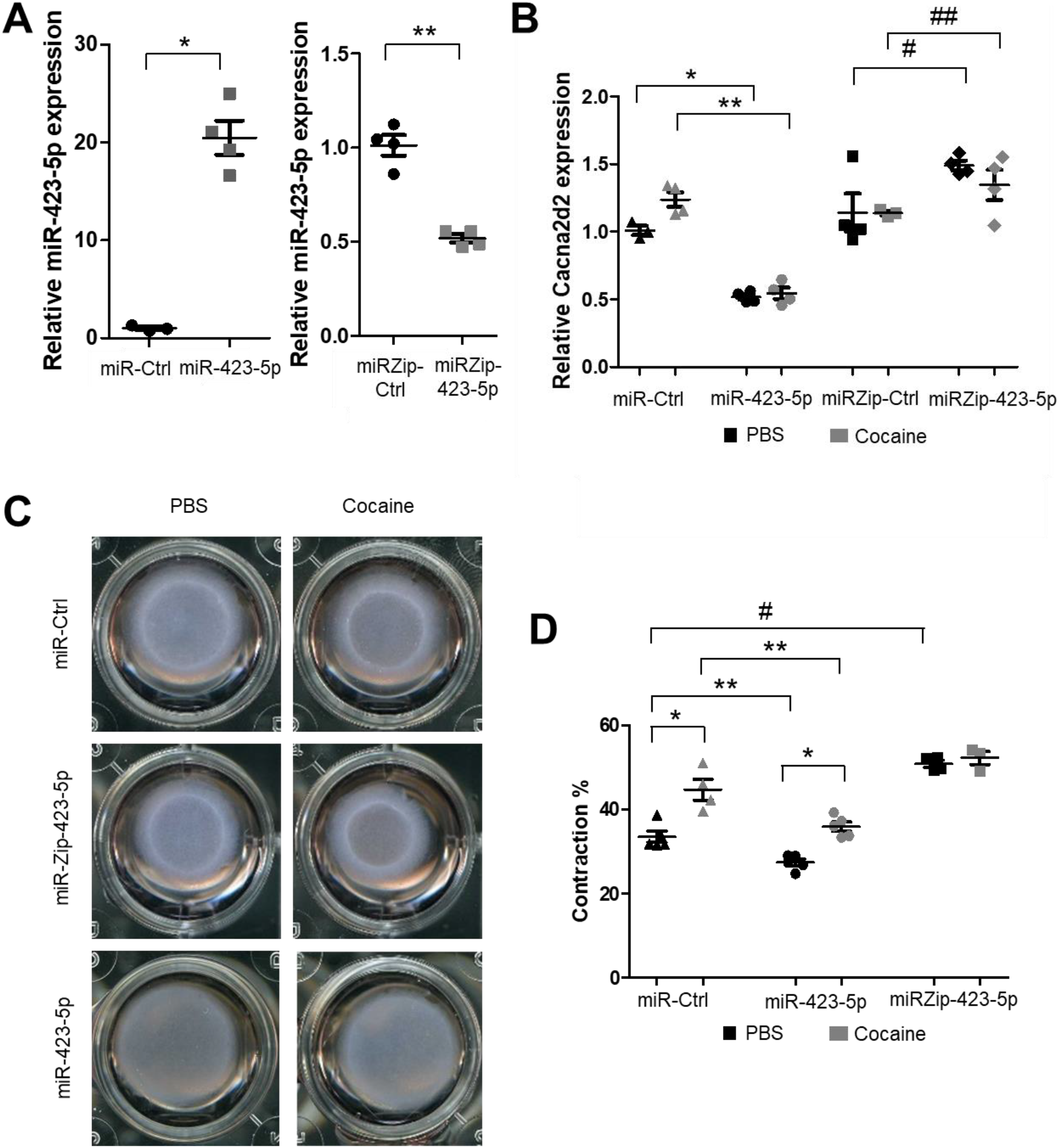
The miR-423-5p-Cacna2d2 axis mediates cocaine-induced SMC contraction. SMCs were transduced with lentiviral vectors encoding control miRNA (miR-Ctr), miR-423-5p, Control miRNA antagonist (miRZip-Ctr) or the miR-423-5p antagonist (miRZip-423-5p). (**A**) qRT-PCR analysis shows that miR-423-5p expression was increased in the miR-423-5p transduced cells and decreased in miR-Zip-423-5p transduced cells (*p<0.05 vs. miR-Ctr;**p<0.05 vs. miRZip-Ctr). (**B**) MiR-423-5p overexpression led to a significant decrease in Cacna2d2 expression both in the absence (PBS) or presence of cocaine (p<0.05 compared to miR-Ctr). Conversely, silencing of miR-423-5p by miRZip-423-5p treatment led to increased Cacna2d2 expression in the presence or absence of cocaine (*p< 0.05 vs. miR-Ctr + PBS; **p<0.05 vs. miR-Ctr + cocaine; #p<0.05 vs. miRZip-Ctr + PBS; ##p<0.05 vs. miRZip-Ctr + cocaine). (**C**) Aortic SMCs were treated with lentiviral vectors encoding miR-423-5p, miR-Ctr, miRZip-423-5p or miRZip-Ctr, and the transduced SMCs were seeded on collagen gel with or without treatment of cocaine. Representative images of the collagen contractility assay show miR-423-5p abrogates, whereas miRZip-423-5p potentiates cocaine-induced SMC contraction. (**D**) Quantification of collagen gel contractility assays confirm that cocaine-induced SMC contraction is mediated, at least in part, by miR-423-5p (*p<0.05 vs. Saline; **p<0.05 vs. miR-Ctr; #p<0.05 vs. miR-Ctr).

To determine if the increased SMC contraction in response to cocaine and CM was mediated via the miR-423-5p—Cacna2d2 pathway, collagen gel contraction assays were performed in SMCs transduced with lentivirus encoding miR-423-5p or the miR-423-5p antagomir – miRZip-423-5p. Overexpression of miR-423-5p blocked cocaine-induced SMC contraction compared to the control miRNA (miR-Ctrl) treated cells. Interestingly, miR-423-5p overexpression also led to a decrease in contractility in the absence of cocaine treatment (**Fig 3C and D**). By contrast, silencing of miR-423-5p expression (miR-Zip-423-5p transduced SMCs) showed increased SMC contraction, irrespective of cocaine treatment, compared to miR-Ctrl transduced SMCs. These results show that the modulation of the cocaine responsive miRNA – miR-423-5p – altered SMC contractility with the overexpression of miR-423-5p suppressing SMC contraction while, conversely, the silencing of miR-423-5p (miRZip-423-5p treatment) promoted SMC contraction.

Each miRNA has the potential to target multiple mRNAs^25^. Therefore, we sought to confirm that the effect of miR-423-5p on SMC contraction was regulated through Cacna2d2. To that end, Cacna2d2 expression was silenced in SMCs (Cacna2d2 siRNA) both in the presence and absence of cocaine (**Fig 4A**). Cacna2d2 silencing significantly abrogated the contractility of the SMCs in the Collagen matrix contractility assay upon cocaine exposure, compared to the control siRNA treated cells (**Fig 4B-C**). To further confirm the role of [Ca^2+^]_i_ in cocaine-induced SMC contraction, SMCs were treated with the L-type voltage gated Ca^2+^ channel blocker Nimodipine (NM) in the presence or absence of cocaine. Treatment with NM resulted in decreased [Ca^2+^]_i_ (**Supplemental Figure 1**) compared to control treatment. Importantly, NM blocked the cocaine-induced SMC contraction (**Fig. 4D and E**) showing that the SMC contraction was mediated through altered intracellular Ca^2+^ levels. Remarkably, the effects of NM on cocaine -induced [Ca^2+^]_i_ and SMC contraction were similar to that seen with Cacna2d2 silencing– either mediated by the overexpression of miR-423-5p (**Fig. 3C and D**) or the siRNA-mediated silencing of Cacna2d2 (**Fig. 4B and C**). These data confirm that cocaine-induced SMC contraction was mediated via the miR-423— Cacna2d2—[Ca^2+^]_i_ axis.

**Figure 4.**
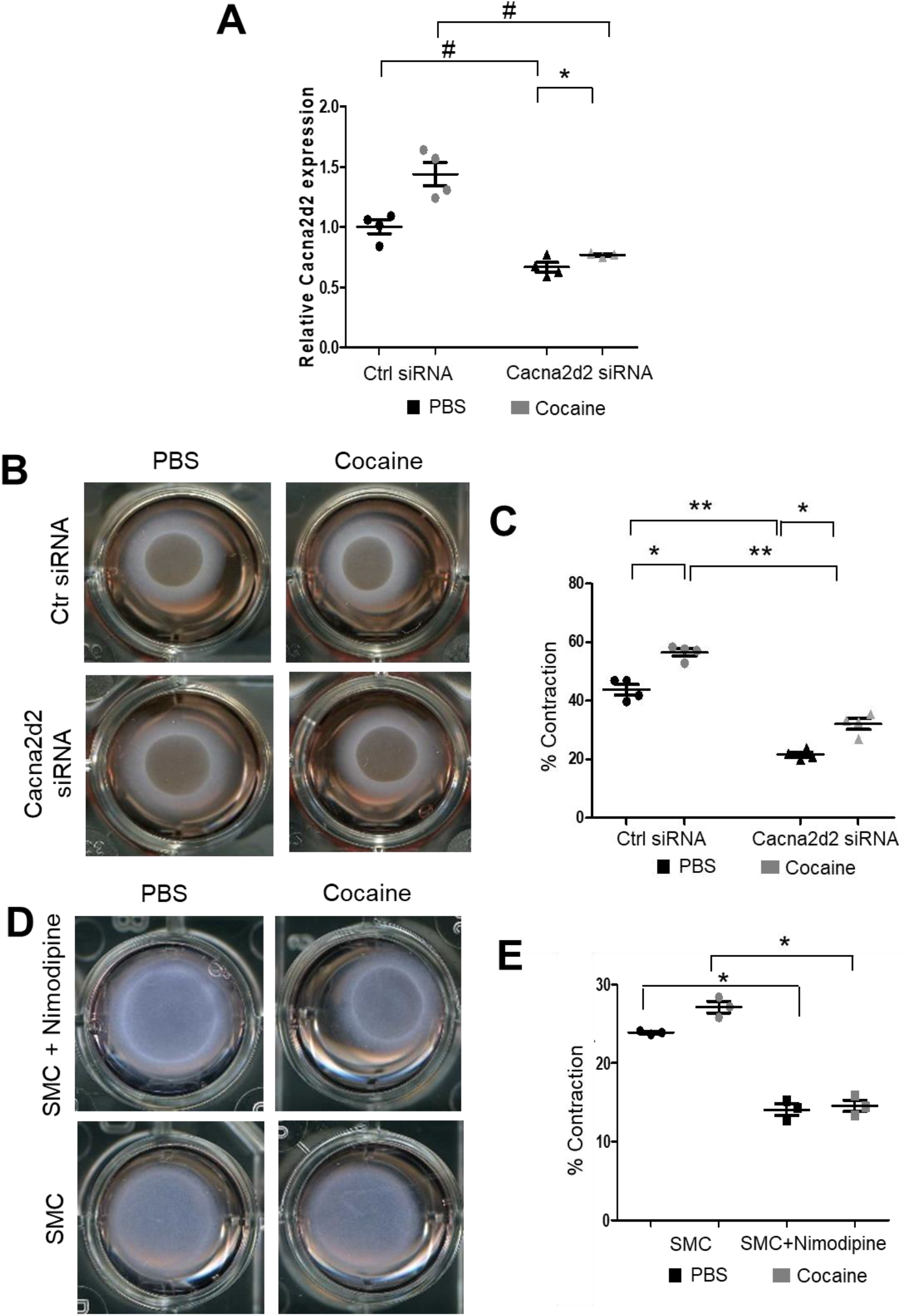
Cacna2d2 and intracellular calcium concentration mediate cocaine-induced SMC contraction. (**A**) Cacna2d2 siRNA decreased Cacna2d2 mRNA expression in SMCs compared with Ctr siRNA in the absence or presence of cocaine (*p<0.05 vs. cocaine; ** p< 0.01 vs. cocaine; #p<0.05 vs. Ctr siRNA). (**B**) Representative images of the Collagen contractility assay in SMCs in which Cacna2d2 was silenced in the absence or presence of cocaine show that the silencing of Cacna2d2 in SMCs leads to a decrease in SMC contractility compared to Ctr siRNA. (**C**) The effects of Cacna2d2 silencing on SMC contractility was measured by image J analysis of the collagen gel contractility assay (*p<0.05 vs. saline treatment; **p<0.01 vs. Ctr siRNA). (**D**) Treatment of SMCs with the L-type calcium channel antagonist Nimodipine (NM) reduces SMC contractility (compared to PBS treatment) and abrogates the effect of cocaine on SMC contractility. (**E**) Quantification of collagen gel contraction images show NM inhibited SMC contraction compared with PBS treatment and abrogated SMC contraction induced by cocaine (*p<0.05 vs PBS).

### 4. miR-423-5p ameliorated Cocaine-induced Increases in BP and Aortic Stiffness *in vivo*

In primary SMCs, cocaine exposure resulted in an elevated [Ca^2+^]_i_ and induced SMCs contraction by modulating the miR-423-5p — Cacna2d2 pathway. To determine if the miR-423-5p — Cacna2d2 axis contributes to the elevated BP and aortic stiffness seen in the mouse model of cocaine use/abuse, mice were injected with lentivirus that overexpress miR-423-5p under the control of a SMC-specific *Sm22*α promoter or an empty vector control lentivirus. These mice were then exposed to daily intraperitoneal injection of cocaine over the course of 10 days. Aortas harvested from control transduced and cocaine-treated mice showed increased Cacna2d2 expression compared to the saline treatment. MiR-423-5p lentivirus transduction reduced Cacna2d2 expression in both cocaine and saline treated mice aortas compared to the control lentiviral transduction (**Supplemental Fig. 2**). BP was measured throughout the course of the experiment with both systolic and diastolic BP showing a progressive increase over the course of the experiment in the control transduced mice following cocaine treatment compared to the saline treatment **(Fig. 5A and B**). However, pre-treatment with the miR-423-5p overexpression vector partially abrogated the cocaine induced increase in both systolic and diastolic BP compared to the control treatment **(Fig. 5A and B**). Aortic stiffness was measured at day 0 and then 2 days following the last cocaine treatment by pulse wave velocity (PWV). Similar to the results of the BP analysis, control transduced mice showed increased aortic stiffness after cocaine exposure when compared with saline treatment. Importantly, transduction of the mice with miR-423-5p expressed from the Sm22α promoter was able to ameliorate the increase in PWV seen with cocaine treatment (**Fig. 5C)**. Collectively, these results indicate that the cocaine-induced increases in BP and aortic stiffness can be, at least partially, abrogated by miR-423-5p overexpression, providing *in vivo* evidence that the miR-423-5p – Cacna2d2 – [Ca^2+^]_i_ pathway plays an important role in mediating the effects of cocaine on BP and aortic stiffness.

**Figure 5.**
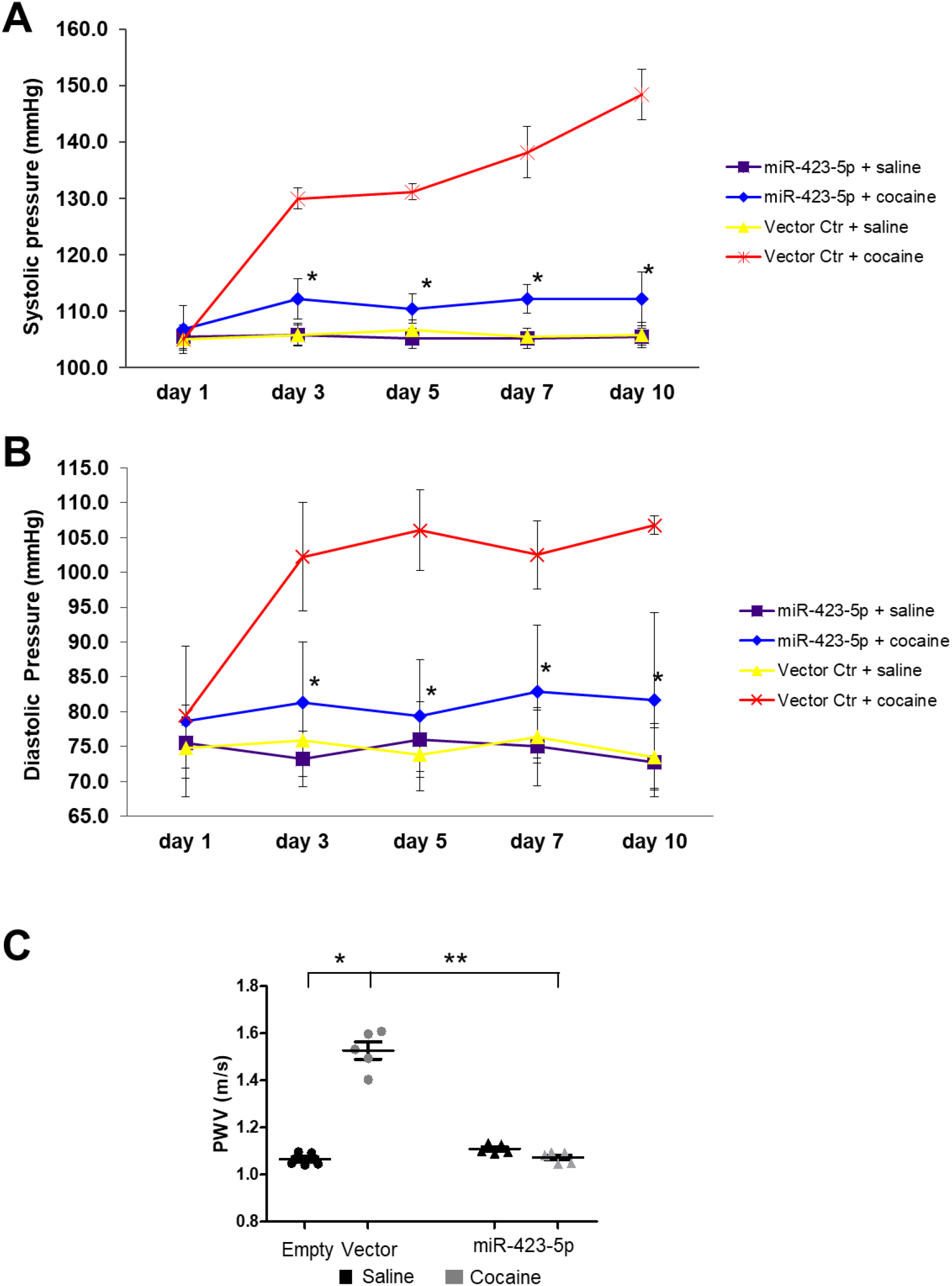
SMC-specific miR-423-5p ameliorated Cocaine-induced increase BP and aortic stiffness in mice. Mice were injected with lentivirus encoding miR-423-5p from the smooth muscle-specific promoter (Sm22α) or vector control before exposure to cocaine or PBS for 10 consecutive days. Pretreatment with miR-423-5p resulted in a partially reduction in the cocaine-induced (**A**) systolic and (**B**) Diastolic BP elevation. (*: p<0.05 vs. miR-Ctr + cocaine). (**C**) Aortic stiffness was measured by pulse wave velocity (PWV) two days following the final treatment with cocaine. Pretreatment of the mice with the miR-423-5p expressing vector abrogates cocaine-induced elevation in aortic stiffness. (*p<0.05 vs. miR-Ctr + cocaine; **p<0.01 vs. miR-423-5P + cocaine).

## DISCUSSION

The CV consequences of cocaine exposure have been well documented (recently reviewed in^52^). These include both short term (acute) effects, including acute chest pain, acute coronary syndrome (ACS), hemorrhagic and ischemic stroke, and cardiac arrhythmias^53^, as well as long-term consequences, including HTN, aortic stiffness, increased left ventricular mass, and bradycardia^9-11^. Cocaine acts through multiple mechanisms to exert it’s effects on the CV system^54,55^. Cocaine stimulates the sympathetic nervous system by increasing the sensitivity of adrenergic nerve terminals to norepinephrine and inhibiting catecholamine reuptake at sympathetic nerve endings^12^. These sympathomimetic effects led to the stimulation of cardiomyocyte adrenergic receptors resulting in increased heart rate, BP, and vasoconstriction that, in turn, increases myocardial oxygen demand. The sympathomimetic effects can cause elevated levels of the vasoconstrictor protein endothelin-1^56^, increased ROS levels^30^, inhibition of nitric oxide (NO) synthase^19^, impaired acetylcholine-induced vasorelaxation^57^, and the dysregulation of intracellular calcium levels^58^. Although the sympathomimetic mechanisms contribute to CV disorders, they can’t explain the full extent and the diversity of cocaine-induced CV phenotypes.

Accumulating experimental and clinical evidence supports a key role for miRNAs in regulating a wide variety of cellular processes, including the response to cocaine exposure ^59-62^. For example, miR-212 was found to be elevated in the dorsal striatum following cocaine exposure regulating compulsive-like cocaine self-administration^31^. This effect was mediated through the modulation of cAMP response element binding protein (CREB) signaling. MiR-495 was shown to be down regulated in response to an acute exposure to cocaine leading to an upregulation of the miR-495 target genes – brain-derived neurotropic factor (BDNF), Calcium/Calmodulin Dependent Protein Kinase II Alpha (CaMKIIα), and Activity Regulated Cytoskeleton Associated Protein (Arc)^63^. The overexpression of miR-124 or let-7 in the Nucleus accumbens attenuated cocaine-induced conditioned place preference in rats^64^. These cocaine-responsive miRNA regulatory pathways have all been studied in the central nervous system. Far less is known about the role of miRNAs in mediating the cocaine effects in the CV system. Recently, we demonstrated that miR-30c-5p was upregulated in the aortas of cocaine exposed mice leading to the downregulation of Me1 and an increase in ROS^30^. This elevation of ROS contributed, in part, to the increased BP and aortic stiffness seen in cocaine or CM exposed mice. This cocaine-induced elevation in BP and aortic stiffness could be partially abrogated by pretreating the cocaine exposed mice with a miR-30c-5p antagonist. In an effort to uncover additional miRNAs that could mediate the cocaine effects in the CV system, we focused on the analysis of miR-423-5p, a miRNA that has been implicated in heart failure, coronary artery disease, pericardial effusion after cardiac surgery^34^, as well as ischemia/reperfusion and the induction of apoptosis^65^. We found that miR-423-5p was markedly decreased in aortas from mice treated with a 10-day course of cocaine. Furthermore, miR-423-5p expression was anti-correlated with the calcium voltage-gated channel component Cacna2d2, a predicted target of miR-423-5p. Luciferase reporter assays showed that Cacna2d2 was a direct target of miR-423-5p. By modulating Cacna2d2 expression, miR-423-5p was found to control [Ca^2+^]_i_ and, in turn, SMC contractility. It is well established that the regulation of [Ca^2+^]_i_ levels plays a critical role in regulating SMC contractility and the development of myocardial ischemia, infarction, HTN, and arrhythmia^18,49,58,66^. L-type Ca_v_1.2 channels (LTCCs) are the principal channels involved in mediating the influx of Ca^2+^ and the regulation of myogenic tone^49-51^ Overexpression of miR-423-5p resulted in reduced SMC contractility in the presence of cocaine. Conversely, the silencing of miR-423-5p expression resulted in increased SMC contraction. Similar to miR-423-5p overexpression, Cacna2d2 silencing using small interfering RNA resulted in decreased SMC contraction. Importantly, the overexpression of miR-423-5p from a SMC specific promoter (*Sm22*α) was able to partially abrogate the elevation of systolic and diastolic BP and aortic stiffness observed in a mouse model of cocaine use/abuse. These results showed that the miR-423-5p—Cacna2d2—intracellular calcium concentration ([Ca^2+^]_i_) pathway serves as an important regulator of BP and aortic stiffness in response to cocaine.

Voltage-gated calcium channels (VGCCs) are protein complexes composed of a main, pore-forming α _1_ subunit and auxiliary α 2δ and β subunits with these auxiliary subunits thought to modulate the biophysical properties of the channel and participate in the trafficking and surface expression of the calcium channel^67^. Previous studies have shown that Cacna2d2 was a direct target of another miRNA, miR-1231^68^. Zhang and colleagues (2017) showed that miR-1231 was overexpressed in both human hearts following MI and in the hearts of rats subjected to an experimental model of MI^68^. Suppression of miR-1231 expression in rat hearts abrogated arrhythmias in the MI model by modulating Cacna2d2 levels. These results indicate that Cacna2d2 may serve as the intersection point between several miRNA pathways in CV disease facilitating the modulation of this key Ca^2+^ channel component in response to different environmental challenges. The effects of modulating Cacna2d2 activity are not restricted to CV tissue. In the nervous system, enhanced expression of the α2δ-2 subunit increased CaV2 channel density at the presynaptic active zone leading to increased cytosolic-free calcium concentrations^69^. The ducky (du) mutation of Cacna2d2 in mice showed a severe phenotype characterized by cerebellar ataxia, epilepsy, reduced body weight, and premature death^70^. Cerebellar Purkinje cells of du mice had 35% smaller whole-cell Ca^2+^ currents mediated by P-type (Ca_v_2.1) VGCC calcium channels and abnormal morphology of their dendritic trees^71^. Furthermore, the α_2_δ-2 subunit regulates the Ca^2+^ current amplitude and impacts the gating of L-type (Ca_v_1.3) VGCC calcium channels in du mice^72^. Cacna2d2 null mice showed growth retardation, cerebellar degeneration, elevated susceptibility to seizures, reduced life span, and cardiac abnormalities, including alterations in heart rate (bradycardia).

MiRNAs have the potential to silence multiple targets increasing the regulatory capacity of these molecules. MiR-423-5p has been shown to interact with multiple target genes. For example, Zhang et al (2021) ^73^ showed that miR-423-5p levels were upregulated in exosomes from the plasma of bicuspid aorta valve (BAV) patients and could regulate TGF-β signaling by decreasing SMAD2 expression. Furthermore, miR-423-5p has been shown to regulate cell apoptosis by enhancing caspase 3/7 activity and impairing mitochondrial functionality by directly targeting Myb-related protein B (MYBL2) expression in a model of hypoxia/reoxygenation^74^. MicroRNA-423-5p has also been shown to target O-GlcNAc transferase to induce apoptosis in cardiomyocytes in response to oxidative stress (H_2_O_2_ treatment)^34^. These results show that miR-423-5p plays multiple roles in CV phenotypes in response to different environmental cues. Cocaine exposure has multifactorial effects on the CV system by engaging multiple miRNA-mRNA pathways that target different aspects of CV cell health and functionality. We have already uncovered important roles for the miR-30c-5p – Me1 – ROS pathway and, in this work, the miR-423-5p – Cacna2d2 – [Ca^2+^]_i_ in the CV response to cocaine exposure. The therapeutic modulation of these pathways may prove to be beneficial in the treatment of cocaine-induced CV phenotypes, including HTN and vascular senescence (aortic stiffness).

## ACKNOWLEDGEMENTS

We would like to thank the Animal Care Team from the Division of Veterinary Resources (DVR) from the University of Miami for their outstanding work.

## HIGHLIGHTS

- MicroRNAs and their pathways are involved in the cardiovascular effects of cocaine.
- MiR-423-5p and its target Cacna2d2 regulate intracellular calcium concentration and smooth muscle cells contractility, which, in turn regulates blood pressure.
- Modulation of the miR-423-5p—Cacna2d2—Calcium transport pathway may represent a novel therapeutic strategy to improve cocaine-induced hypertension and aortic stiffness.

